# Axons-on-a-Chip for Mimicking Non-Disruptive Diffuse Axonal Injury underlying Traumatic Brain Injury

**DOI:** 10.1101/2021.05.06.442958

**Authors:** Wei Li, Haofei Wang, Xiaorong Pan, Dejan Gagoski, Nela Durisic, Zhiyong Li, Chun-Xia Zhao, Tong Wang

## Abstract

Diffuse axonal injury (DAI) is the most severe pathological feature of traumatic brain injury. However, how primary axonal injury is induced by mechanical stress and whether it could be mitigated remain unknown, largely due to the resolution limits of medical imaging approaches. Here we established an Axon-on-a-Chip (AoC) model for mimicking DAI and investigating its early cellular responses. By integrating computational fluid dynamics and microfluidic techniques, DAI was observed for the first time during mechanical stress, and a clear correlation between stress intensity and severity of DAI was elucidated. This AoC was further used to investigate the dynamic intracellular changes occurring simultaneously with stress, and identified delayed local Ca^2+^ surges escorted rapid disruption of periodic axonal cytoskeleton during the early stage of DAI. Compatible with high-resolution live-microscopy, this model hereby provides a versatile system to identify early mechanisms underlying DAI, offering a platform for screening effective treatments to alleviate brain injuries.

## Introduction

In severe head injuries, rapid accelerations-decelerations of the brain tissue in the rigid skull cavity cause diffuse axonal injury (DAI) to the long axons connecting brain regions and hemispheres ^1^. DAI, as the most serious form of traumatic brain injury (TBI), often leads to death or long-term disability for patients. Pathology studies have shown that DAI is characterized by multifocal lesions scattering in long axons within the white matter, most commonly in internal capsule, corpus callosum, cerebellar peduncle, brain stem and gray-white boarder of the cerebrum ^2^. These lesions result in devastating neurological symptoms such as coma, unconsciousness, paralysis and cognitive deficits, and is therefore a poor prognostic indicator ^3^. Subcellularly, DAI is caused by the shearing of the axonal structure due to the differential force-driven movement between the neuronal cell body and its axon, which exhibit the greatest density difference ^1,2^. Indeed, with a typical width to length ratio more than 1 million, the long but extremely thin projecting axons of central neurons are particularly vulnerable to mechanical impacts from vertical direction. Relative motion and friction between adjacent matrix and cells could exert a joint of compression and shear force to the axonal plasma membrane ^4^. Such a force leads to an influx of extracellular calcium (Ca^2+^), which, as observed *in vitro*, appears within several minutes and initiates different signaling cascades for further axonal degeneration ^5,6^. Besides the surging of intracellular Ca^2+^ waves, formation of multiple focal axonal swellings (FAS), retraction balls and halt of axonal trafficking cargoes were also noticed along the axon shafts, signifying the activation of neurodegeneration ^7–11^.

Limited by the resolution of current noninvasive medical imaging techniques, these injury-induced instant intra-axolemma changes could not be detected in either patients or animal models ^12^. To address this critical issue, various *in vitro* models have been established to study the earliest cellular cascades triggering injury. Among multiple *in vitro* models, the Polydimethylsiloxane (PDMS) microfluidic devices have recently been developed to investigate the axonal injury because of their advantages such as good biocompatibility, optical transparency and easy fabrication with low cost ^13,14^. Compared to traditional mix-cultured neurons, it also offers significant advantages, including precisely controlled lesion to multiple axons and compatibility with high-resolution cell-biological assays such as live-imaging, electronic and super-resolutions microscopy ^15,16^. Cellular responses to mechanical insults have been studied using two types of microfluidic-based models, complete axotomy by severing of multiple axons ^13,17,18^, or incomplete axonal damage by either axial over-stretching ubiquitously of the whole neuronal cell body ^19,20^ or focal crushing of short axonal segments ^21^. However, none of these *in vitro* models fully represent the typical clinical DAI, which comprises of wide-spread micropathological changes along axons caused by force from multiple directions, as observed in severe close-skull TBI ^22^. Therefore, a new reliable *in vitro* model is urgently needed for DAI research, to observe the subcellular response immediately after, or even during the process of mechanical stress.

In this study, we developed a microfluidic-based Axons-on-a-Chip (AoC) platform to model DAI induced by mechanical stress in living neurons. To achieve the precise control of damaging stress, we first conducted numerical simulation using computational fluid dynamics (CFD) to calculate the amplitude of mechanical impact on axonal surface. By simulating the stress distribution on axonal surface, we established the relationship between the injecting fluid and the mechanical stress exerted onto the axonal surface. Then, we conducted microfluid-induced axonal damage in live neurons to investigate the extent of DAI caused by various injecting flow rates and directions using live-imaging confocal microscopy, and found that DAI severity is positively correlated with both relative angle and injecting flow rate. Finally, using Ca^2+^ imaging and time-lapse super-resolution microscopy, the instantaneous subcellular responses during the mechanical impact were investigated. Our study for the first time establishes an *in vitro* AoC platform to precisely manipulate the degree of DAI, and explores its instant subcellular responses, and hereby advances the insights into new mechanism underlying TBI.

## Results

### Design of microfluidic AoC device to mimic the mechanical force-induced DAI

DAI represents the primary cellular damage caused by the mechanical impacts and happens shortly after the severe brain trauma. However, lesions signifying the DAI in the white matter can only be detected by non-invasive medical imaging approaches immediately afterwards, when the most severe damage and degenerative fate of the neurons are determined ^3^. Using cultured primary neurons, *in vitro* studies identified that rapid cellular changes are induced by the mechanical impacts seconds to minutes after initiating cascades that lead to irreversible diffuse axonal damage ^23–25^. Taking advantage of the extreme length of the projecting axons, microfluidic-based AoC devices were designed to study the degenerative or regenerative responses following the disruptive axonal severing (axotomy) ^4,17,18^.

To further observe the instantaneous effects of non-disruptive DAI, the AoC platform in this study was designed. One middle injury chamber and two side chambers (reservoirs) were interconnected by arrays of 80 microchannels. Dimension of each component was shown in the schematic illustrations in Fig. 1A. The middle injury chamber was designed with a height of 105 μm and a width of 1000 μm to allow medium perfusion. The two ends of middle chamber was punched to create inlet and outlet, which were connected to a syringe pump allowing fluid flow at a controlled rate and duration. The two side chambers (5 mm × 5 mm) located on the opposite side of the middle injury chamber were punched to create open reservoirs for neuron inoculation and medium replacement. The middle chamber was connected to the side reservoirs via 80 microchannels with the dimension of 5 μm (height) ×12 μm (width) × 200 μm (length) and a in-between gap of 50 μm. The small dimension of these microchannels block the migration of neuron cell bodies while allowing the passage of axons only. This PDMS-device was next permanently bonded to the surface of 0.17 μm glass-bottom dish, which is compatible with time-lapse inverted confocal or super-resolution microscopy. Then rat hippocampal neurons were seeded into the two opposing soma chambers and cultured *in vitro* for 7 days (DIV7). After axons with various orientation extended into the middle chamber, microfluidic jets of culture medium were pumped into the middle chamber to exert various extents of mechanical force onto the crossing axons, which could cause the non-disruptive DAI as well as the disruptive axotomy (Fig. 1B). To capture the real-time subcellular changes following the fluid-induced injury, mechanical stress applied on axons is precisely controlled by adjusting the flow injected via inlet tubings (Fig. 1C), while instantaneous effects of non-disruptive mechanical damages on diffuse axonal sites were observed by inverted live-imaging microscopy (Fig. 1D-F).

**Figure 1.**
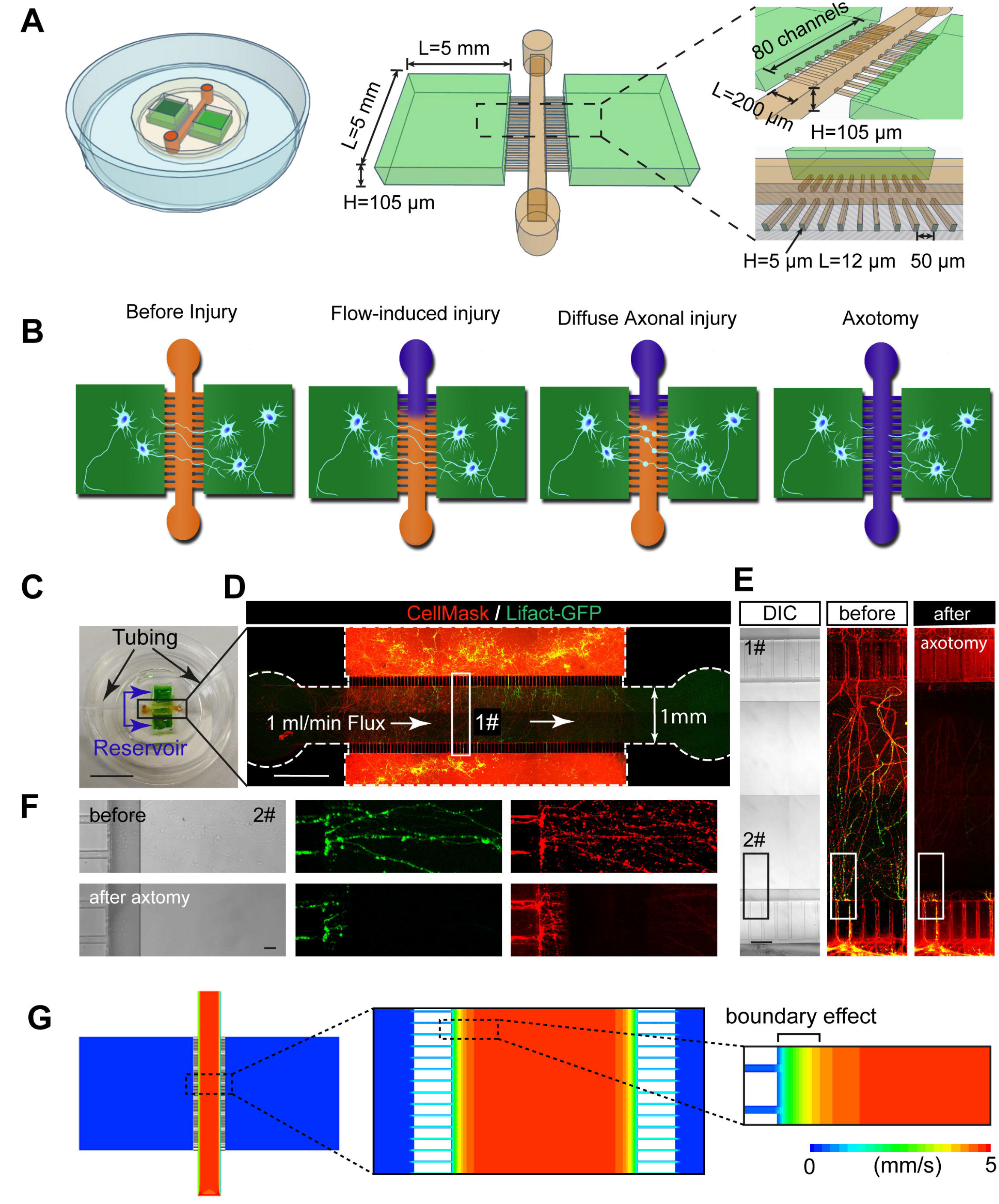
Design of the Axon-on-a-Chip (AoC) device to induce mechanical injury to axons of live hippocampal neurons. **(A)** 3D representation of the microfluidic device showing dimensions of different compartments. **(B)** Schematic cartoons showing the procedure of microfluidic flow-induced non-disruptive diffuse axonal injury (DAI) and axotomy, which were induced exclusively in the axonal segments using different flow rates. **(C)** Photo of the fabricated AoC device attached to the glass-bottom dish, with reservoirs (green) and central injury channel (yellow) labelled with different colours. Bar = 1 cm. **(D)** Live-imaging microscopic graphs showing the axon microchannels and the central injury channel, with images of axons dual-labelled with Lifact-GFP (Cytoskeleton marker) and CellMask (Plasma membrane) in the central channel were captured before and after the 1 mL/min-flow injection. Bar = 1 mm. **(E)** Amplification of the boxed region (1#) in **(D)**, showing axons before and after the 1mL/min flux induced disruptive axonal injury (axotomy) in the cross section of the central injury chamber. Bar =100 μm. **(F)** The region (2#) proximate to axon channels in **(E)** were shown before and after the axotomy. Dual-labelled axons were fully disrupted by the 1 mL/min flux, with remaining stubs near the edge of the device indicated with arrowheads. Bar = 20 μm. **(G)** Simulated velocity distribution inside the microfluidic device showing the decreased flow velocity near the boundary of the device with the injection flow rate of 1000 μL/min.

To examine whether the jet flow can induce mechanical damages to the axons, we first injected the culture medium at a very high speed of 1000 μL/min, which equals to 9.13 Pascal (Pa) in pressure intensity. We found that axotomy and removal of axonal segments occurred in the middle chamber following 3 min of the jet flow (Fig. 1D-E). Interestingly, in highly amplified images, we also found the severing of the axon was restricted to the proximal of PDMS boundaries, leaving the stub ends and axonal shafts intact near the boundary and in the microfluidic grooves (Fig. 1F), indicating the fluid-induced stress is significantly reduced towards the solid boundary. This phenomenon can be explained by the boundary effect of fluid dynamics that the flow velocity rapidly reduces near the boundary with the injecting flow rate of 1000 μL/min (Fig. 1G). Accordingly, this also reminds us that by altering the flow rate of medium injection, it is possible to induce various levels of cellular injury responses at single axon level. Collectively, these results suggest that the AoC device could mimic the mechanical stress-induced axonal damage, with the potential to detect cellular responses under different magnitudes of stress

### Numerical simulation of the mechanical stress by microfluidic jet injection

We next sought to simulate the stress distribution within the middle injury chamber using CFD ^26^, thus precisely controlling the extent of DAI (Fig. 2A). The stress that jet fluids exerted on axons includes shear stress and normal stress, which are parallel or vertical to the axon surface, respectively. As the normal stress is the dominant cause of DAI ^21^, we adapted the pressure contour to qualitatively describe the mechanical effects of injecting flow and the average normal stress (average normal force per acting area) on the crossing axon. To investigate the effect of the reservoirs on axon damage, we first generated and compared the shear stress distribution on the bottom surface of the AoC device with (Fig. 2B-C, top panels) or without (Fig. 2B-C, bottom panels) considering the dimensions of reservoirs. There was no obvious difference between these stress contours. Eight different water injection flow rates were simulated and compared, no obvious difference among the normal stress on axon was observed (Fig. 2D), indicating that we could neglect the effect of reservoirs on the normal stress exerted on axon surface. It also clearly shows that the normal stress on axon surface increased linearly with the flow rate, so we can use different flow rates to simulate different mechanical stress to investigate axon injury. Furthermore, we also examined whether the location of axons crossing the injury chamber affects the normal stress induced by flow injection, we calculated the normal stress on one or eighty-one crossing axons in the central injury chamber, respectively. To observe the normal stress exerted on axon surface quantitatively, we compared the pressure contour in three different flow rates on the same axon location (Fig. 2E). We found that no obvious difference of pressure distribution on the axon in the same location was observed between one and eighty-one axons. We next analyzed the average normal stress on each axon with eighty-one axons inside the injury chamber in eight different flow rates (Fig. 2F). It clearly showed that the normal stress exerted on each axon was almost the same using the same flow rate, verifying that the axon location did not have any significant effect on the flow-induced normal stress. We further compared the average normal stress distribution between one and eighty-one axons inside the central chamber (Fig. 2G). The average normal stress on each axon does not change with either the location or the number of the crossing axons, and there is a linear correlation between the flow rate of the injecting flow and its acting normal stress on either one or eighty-one axons (Fig. 2G). These results suggest that a steady flow in the central injury chamber generates a uniform stress distribution over multiple crossing axons, and this applied stress only correlates linearly with the flow rate. We therefore can precisely control the stress on individual axons in the central chamber, by adjusting the flow rate of the injecting fluid.

**Figure 2.**
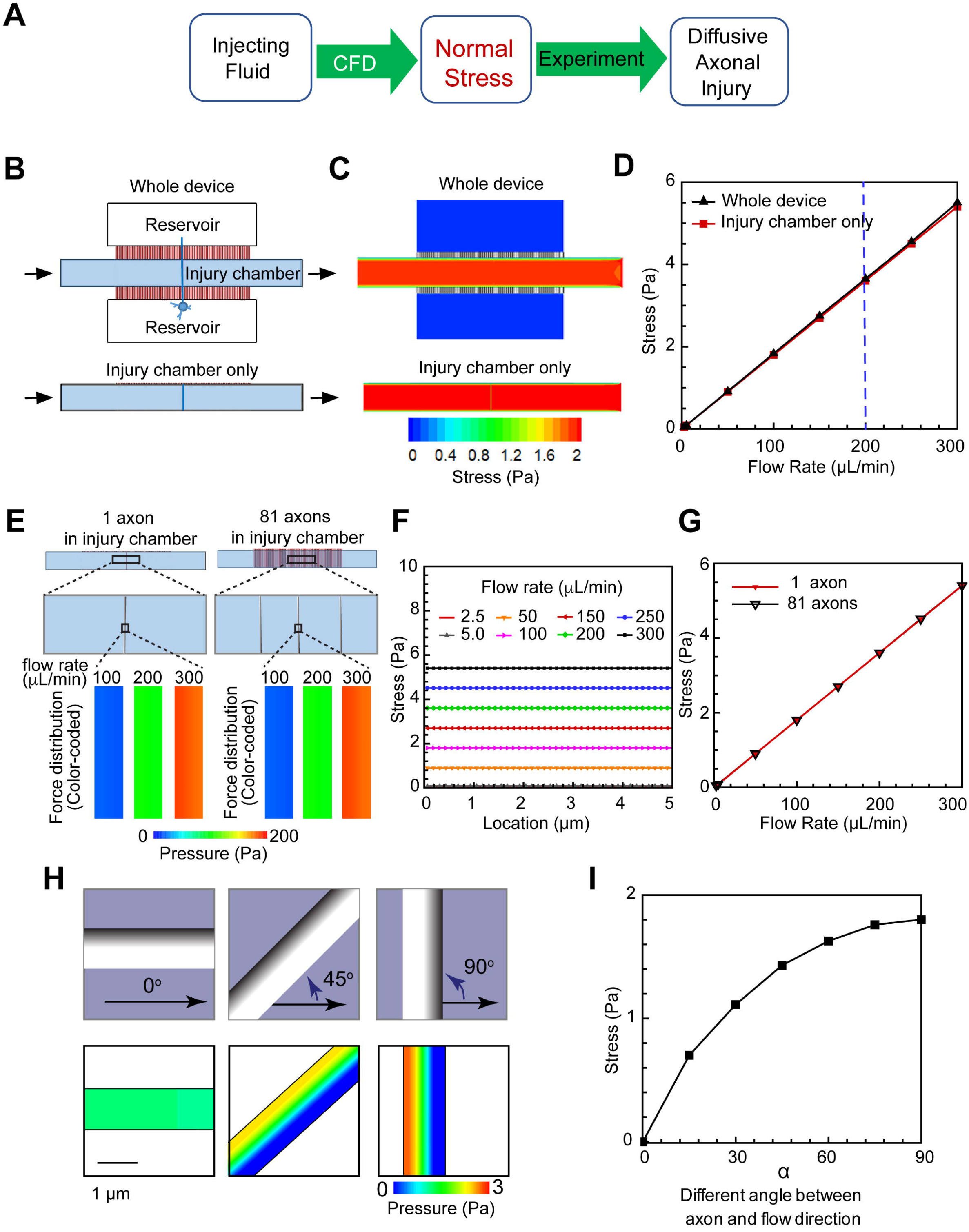
Simulation of the intensity of mechanical stress caused by the microfluidic injection in the AoC. **(A)** Diagrammatic sketch to show the link between computational fluid dynamics (CFD) and experiments using fluid injection. **(B)** Computational domain with and without reservoirs attached. **(C)** Wall shear stress distribution on the bottom surface of AoC device. **(D)** Comparison of the normal stress on single axon with and without reservoirs at eight different flow rates. **(E)** Scheme of the comparison between pressure with one axon and eighty-one axons in injury chamber without reservoirs at three different water injection rates. **(F)** Normal stress distribution on eighty-one axons without reservoirs at eight different water injection rates. **(G)** Comparison of normal stress distribution on eighty-one axons and one axon at different water injection rates. **(H)** Scheme to show the pressure distribution in three different relative angles (α) between flow injecting direction and axon extending orientation. Axons are shown as grey columns in the top panels. **(I)** Impact of angles between axon orientation and flow direction (α) on the normal stress at axon surface with the injection flow rate of 100μL/min.

We also examined the effect of relative angles between the injecting flow direction and crossing axons (α) on the flow-induced normal stress. Similar to Fig. 2E, the pressure contour were adapted to describe the normal stress distribution for three different angles α showing that the pressure difference between the leading edge and trailing edge of the axon increased with the angle (Fig. 2H), resulting in the increase of the average normal stress (Fig. 2I). There is a positive correlation between the normal stress and α values (0°, 15°, 30°, 45°, 60°, 75° and 90°, Fig. 2I) in the range of 0 to 1.8 Pa at the injecting flow rate of 100 μL/min, illustrating minimal normal stress at 0° and maximum at 90°.

To validate these simulation results, we compared these results with those reported in previous studies ^4,27^, where they used either force or stress to describe the mechanical stress induced disruptive or non-disruptive axonal injuries. This mechanical stress, defined in this study as the value of acting surface force per contacting area, is often obtained by either experimental measurement ^21,28^or numerical simulation ^4,27^. For experimental measurement, it is impossible to acquire the comprehensive force distribution along the whole axonal surface due to the limitations of available methods, which often only allow the measurement of an average pressure on discrete contacting loci along the axon ^21,28^. For numerical simulations, the comprehensive force distribution along the surface of the crossing axon could be calculated ^4,27^. We therefore compared the results aquired in this study with those from other numerical studies ^4,27^, Gu et al.^27^ simulated the force using semi-cylinder axons as the model, showing injecting flow rates between 0.5 and 5.5 μL/min could generate forces ranging from 2.0 to 28.0 pN (0.0127 - 0.175 Pa), which is in line with our simulated results of about 0.045 −0.09 Pa for flow rates of 2.5 - 5.0 μL/min. For extreme conditions that could induce complete axotomy ^4,27^, the mechanical force was calculated to be at 485-970 pN, which corresponds to 5.146-10.3 Pa assuming the axon having a diameter of 1 μm.

As we will show later, the AoC system allows a precise control of flow rates over the range of 200 □ 1000 μL/min, corresponding to a normal stress of 1.822 □ 9.1316 Pa. This range of normal stress is capable of inducing both non-disruptive axonal injury and disrupted axotomy, and is in the range of axonal injury ^21,28^ and axotomy ^4,27^. Furthermore, different methods have also been developed to measure non-disruptive mechanical forces of 0 – 7.5 nN (measured using atomic force microscopy) ^21^, 7.63 ± 1.13 nN (stress of 0.27 ± 0.04 nN/μm^2^ and 6 μm-bead) ^29^ and 3.1416 ± 0.754 nN (stress of 0.25 ± 0.06 nN/μm^2^ and 2 μm-microdot) ^21^. These measured forces also fall into the similar range of simulated results derived from this study, which is ~2.826 nN at the flow rate of 200 μL/min. Therefore, the values of CFD-calculated force in the current system are in good agreement with those derived previously using other numerical and experimental methods, demonstrating the current AoC system can be used to model both the non-disruptive axonal injury and disruptive axotomy.

### Formation of focal axonal swellings (FAS) correlates positively with the direction and rate of injecting flow

To examine whether fluid flow could induce DAI of living neurons in the AoC, we first investigated its capability to cause the formation of focal axonal swellings (FAS), which are the early morphological changes of DAI ^30^, and its correlation with the magnitude of mechanical stresses. According to the simulation results, the magnitude of mechanical stress in the injury chamber of AoC is controlled by both the flow rate and the angle between the injecting flow and the axon. Therefore, we should be able to examine the correlation between the extent of DAI and the injecting flow by measuring the diameter and angle of axon before and after the microfluidic injection.

Rat hippocampal neurons transfected with the F-actin marker peptide Lifeact-GFP on DIV 7 were cultured in the AoC microfluidic devices for 3 days, and they were then subjected to flow-induced mechanical injury on DIV 10. Time-lapse confocal images were acquired in the central injury chamber, before, during and after the flow injection (Fig. 3A). We analysed the confocal images by Fiji software, and only the Lifact-GFP channel was selected for analysis. Diameters and minimum angles at multiple sites with equal intervals along the axon shafts were measured at the segments proximal to the channel exits (Fig 3B), at the indicated time points prior to and after the flux-induced injury, respectively. The effect of flow angle on inducing axonal injury was then examined. As shown in Fig 3C, axons sprouted into various orientations crossing the surface of the central injury chamber in the microfluidic device. Results from the CFD modelling predicted that axons with different angles ranging from 0-90° are expected to receive mechanical stresses ranging between 0-1.8 Pa at a flow rate of 100 μL/min. This direction-dependent change of mechanical stress might induce different degrees of axonal injuries. We therefore examined the extent of FAS on different orientated axons induced by the same injecting flow rate (200 μL/min). Using time-lapse confocal microscope, we found that axons with α greater than 45° induce a larger extent of FAS than those with α less than 45° (Fig 3C, right panels). To quantitively describe the extent of FAS, we measured the diameter expansion of axonal segments at 4 time points pior to (Φ_0’_) and following (Φ_5’_, Φ_27’_, Φ_32’_) the 3 min flow injection to reflect the morphological changes of FAS. The value of diameter expansion is an index for DAI magnitude: higher value indicates a more severe degree of DAI. As shown in Fig 3D, axons with α greater than 45° demonstrated more severe lesions after 3 min of flow injection at 200 μL/min. These results indicate that the angle between the axon and injecting flow is positively correlated with the level of DAI, verifying our CFD prediction.

**Figure 3.**
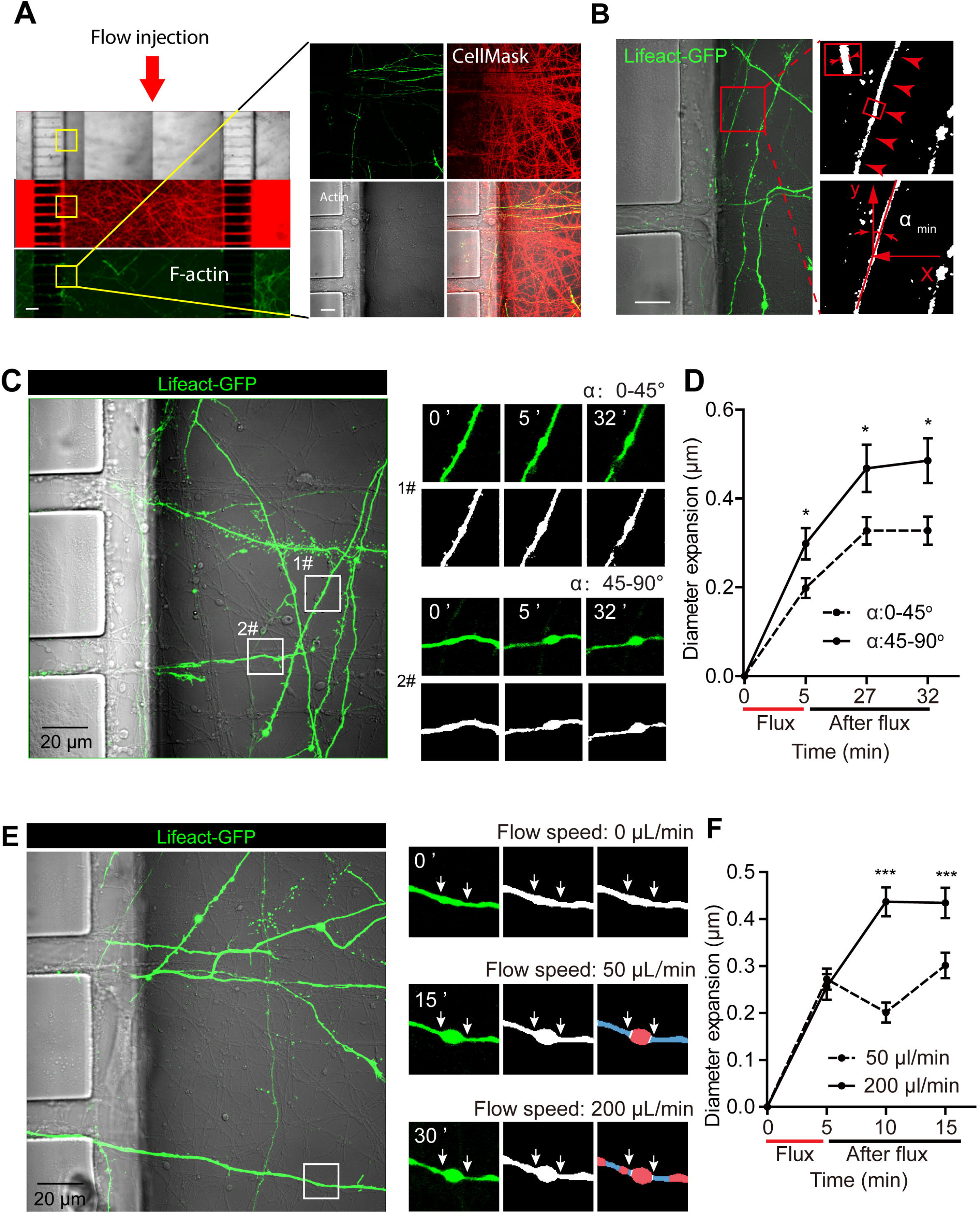
Formation of FAS is correlated with the relative angle and flow rate of injecting microfluidic flow. **(A)** DIV10 rat hippocampal neurons cultured in AoC were transfected with Lifact-GFP on DIV 7 and live-imaged on DIV 10, using LSM710 inverted confocal microscope. Microfluidic flow was injected to the central injury chamber to induce mechanical stress to the crossing axons. Representative confocal images showing the selection of live-imaging window within the injury chamber of AoC device. Injected flow is indicated with red arrow. Amplification of the boxed region was shown in right panels. Bar = 100 μm (left); Bar = 10 μm (right). **(B)** In the terminal chamber, a regions of interest (ROI), which contains at least one axon for analysis. For diameter measurement, five different points with the same distance were chosen, as indicated with red arrows. For tilting angle measurement, the minimum angle (α_min_) was defined as the angle between the axons and that of injected flow, as amplified in the boxed region in the right. Bar=20 μm. **(C)** Representative fields in the terminal chamber with multiple Lifeact-GFP positive axons before the flow injection. Two ROIs were selected on the axons oriented into two groups of different directions: ROI 1# represents the group of axons bearing an α_min_ of 0-45° with that of the injected flow; ROI 2# represents the group of axons bearing an α_min_ of 45-90°. Time-lapse images were shown the morphological changes before and up to 32 min after the 3 min flow injection. For the 1# α_min_: 0-45° and 2# α_min_: 45-90° groups, axonal diameter expansion at 3 indicated time points are shown in the right panels, respectively. **(D)** Quantification of **(C)**, showing the flow-induced diameter expansion induced by 3 min of microfluidic injection. Results are shown as mean±s.e.m, \**p*<0.05, two-tailed student’s *t*-test; angle 0-45°: N = 310, 81, 72, 70 axon segments; angle 45-90°; N = 240, 82, 86, 84 axon segments. **(E)** Time-lapse images showing the morphological changes before and up to 15 min after the 3 min flow injection. For the 50 μL/min and 200 μL/min groups, axonal diameter expansion at 3 indicated time points following the flux are shown in the right panels, respectively. **(F)** Quantification of **(E)**, showing the diameter expansion induced by flow injection. Results are shown as mean±s.e.m, \**p*<0.05, \*\*\**p*<0.0001, two-tailed student’s *t*-test; 50 μL/min group, N=550, 149, 137, 153 axon segments; 200 μL/min group: N= 535, 119, 162, 165 axon segments. Data were from 3 independent cultures.

We next examined the effect of different flow rates on inducing DAI. Time-lapse confocal microscopy was conducted to capture the dynamic change of FAS, before and after three different injecting flow rates of 50 μL/min, 200 μL/min (Fig. 3F) and 1000 μL/min (Fig. 1E-F; see also sMov. 1). Because under the flow rate of 1000 μL/min, most axons were soon completely disrupted (Fig. 1E-F; see also sMov. 1), 1000 μL/min was hereby used as the axotomy condition in our AoC system. Then, by examining the diameter expansion of axon segments before and after the 50 μL/min or 200 μL/min flow, we quantify the extent of FAS, by comparing diameter expansions at different time points, respectively. We found that the 200 μL/min flow induces a more significantly severe DAI than that of 50 μL/min (Fig 3F), suggesting that the intensity of DAI positively correlates with the flow rate of injecting flow, and higher flow rate induces more serious diffuse lesions on axons. These results agree well with our CFD modelling results. Collectively, we found that, in this AoC, different magnitudes of mechanical stress from injecting flow with different angles and flow rate lead to different degrees of DAI.

### Flow-induced rapid primary cellular responses underlying DAI

Due to the fast onsets of primary and secondary injury cascades ^31^, tracing the direct and local axonal changes caused by mechanical stress is technically challenging ^32^. Fundamental questions such as what are the instantaneous cellular responses to the mechanical stress is still unknown. We hereby used the AoC platform to investigate the relevant subcellular alternations simultaneoulsy with the fluid-induced mechanical stress. As the elevated axoplasmic intracellular Ca^2+^ ([Ca^2+^]i) surge is the earliest sign of axonal injury and is pivotal to induce downstream destructive cascades ^31,33,34^, we monitored the instantaneous [Ca^2+^]i upon mechanical stress. To indicate [Ca^2+^]i, hippocampal neurons were transfected with genetically coded calcium sensor GCaMP6-f ^35^ on DIV7, microfluidic injury experiments were performed on DIV10. To monitor the [Ca^2+^]i, high flow speed of 1000 μL/min flow rate was used to induce axotomy (sMov. 1). We found axons were completely disrupted in the axonal chambers at this flow rate (Fig. 4A, bottom; sMov. 1). Interestingly, before the complete disruption of the axon, multiple axonal varicosities were formed with locally elevated [Ca^2+^]i in those expansions (Fig. 4A middle; sMov. 1), which were eventually flushed away and left the stub end with the raised [Ca^2+^]i (Fig. 4A, bottom; sMov. 1). Although the axotomy-induced [Ca^2+^]i increase at the stub end was reported before ^5^, the phenomenon of raised [Ca^2+^]i within the DAI-induced axonal varicosities is less known. Our observation suggests that prior to the complete severing of axon, [Ca^2+^]i elevation were already induced by mechanical stress, leading to diffuse axonal injuries with varicosities. To further characterize [Ca^2+^]i accompanied with varicosity formation along the diffusely injured axon, we used a smaller flow rate of 200 μL/min to specifically induce the non-disruptive axonal injuries (Fig. 4B; sMov. 2). We found that although axons were not disrupted, a significantly increased [Ca^2+^]i could be noticed immediately upon the flux (Fig. 4B, middle panel). For the analysis of this [Ca^2+^]i fluctuation, three time points were chosen: t1 was the timepoint when the flux injection started, t2 was the time that the flux ceased and t3 was chosen after the [Ca^2+^]i surge following the injection, as shown in Fig. 4C. We found that there was a ~1.5-fold [Ca^2+^]i increase from t1 to t2, whereas the [Ca^2+^]i surged to approximately 3.2-fold at t3 (Fig. 4D, E). Simultaneously, the axonal diameter in those varicosities also underwent a moderate increase from t1 to t2, but surged significantly to 3 folds at t3 (Fig. 4F; sMov. 2). Indeed, the extent of focal axonal swelling and [Ca^2+^]i are positively correlated with a Pearson’s coefficient of 0.934±0.02 (Fig. 4G). Our observation of both swellings and [Ca^2+^]i progressed during and following the process of mechanical stress, suggesting a delayed occurrence of DAI in stressed axons.

**Figure 4.**
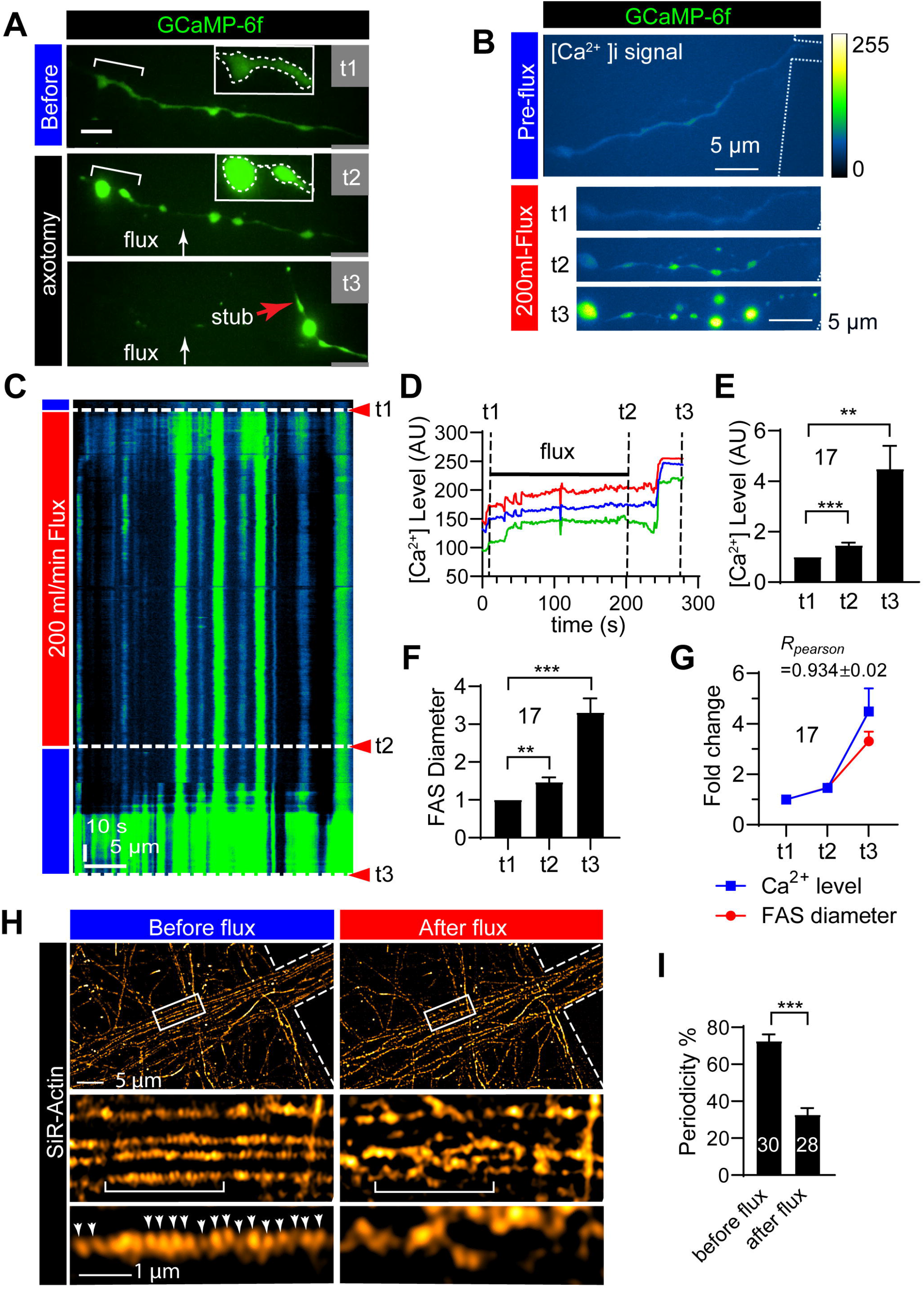
Microfluidic-induced mechanical force initiates rapid intracellular cascades underlying DAI in the AoC. **(A)** Rat hippocampal neurons cultured in microfluidic devices were transfected with GCaMP-6f plasmid on DIV7, then subjected to microfluidic-induced force on DIV10. Time-lapse confocal images were acquired in the central injury chamber, before (t1), during (t2) and after (t3) the microfluidic flow injection. At the flow rate of 1000 μL/min, axotomy was observed in most of the axons. Representative time-lapse image showing the disruptive process of transfected axons is shown, with [Ca^2+^]i level indicated by the fluorescent level of GCaMP-6f. Bar=5 μm. **(B)** At the flow rate of 200 μL/min, non-disruptive axonal damages with diffuse focal swellings were observed along axons. Representative time-lapse image showing the microfluidic-induced injury process along axons, with [Ca^2+^]i level indicated with fluorescent level of GCaMP-6f. Bar=5 μm. **(C)** Kymograph showing [Ca^2+^]i level of an axon exposed to 200 μL/min flow-induced mechanical force, with t1/t2/t3 indicated with arrows. xBar = 5 μm; yBar = 10s. **(D)** Line profiles of [Ca^2+^]i at 3 different FAS sites before (t1), during (t2) and after (t3) the 200 μL/min microfluid-induced injury process. **(E)** Quantification of [Ca^2+^]i changes before (t1), during (t2) and after (t3) the flux injection. **(F)** Quantification of the diameter expansion at the FAS sites before (t1), during (t2) and after (t3) the 200 μL/min microfluid-induced injury process. **(G)** Pearson’s coefficient of FAS diameter and [Ca^2+^]i intensity, showing a positive correlation between these two factors. Data are shown as mean±s.e.m. \*\**p*<0.01, \*\*\**p*<0.0001; N=17 FAS sites. Data were from 3 independent cultures. **(H)** Time-lapse SIM of axons before and after the flux-induced injury were shown, with actin rings in MPS labelled with SiR-Actin. Periodic actin rings were indicated with arrowheads. Bar as indicated in the panels. **(I)** Ratio of periodicity were compared before and after the flux-injury. Data are shown as mean±s.e.m. \*\*\**p*<0.0001; N=30 and 28 before and after the flux, respectively. Data were from 3 independent cultures.

As the structural stability of axon is maintained by the membrane associated periodic cytoskeletal structures (MPS) ^16,36^, and we previously found that the disassembly of axonal MPS leads to the rapid FAS and axonal degeneration ^37^. We next investigate whether the axonal MPS is affected by stress-induced DAI. The axonal MPS were first labelled using the F-actin specific dye SiR-Actin ^27^, then time-lapse structure illumination microscopy (SIM) were adopted to resolve MPS along the bundles of axons formed in the injury chamber as previously established ^37^. Then before and after the 200 μL/min fluid-induced injury, time-lapse SIM images were obtained for the same region of interest (ROI). We performed the paired comparation of the periodicity of the axonal MPS before and after the stress and found that the regular arranged structure of MPS was disrupted in the stressed axons (Fig. 4H), leading to significantly reduced periodicity ratio (Fig. 4I). This phenomenon could be detected as early as several minutes following the mechanical stress, suggesting that the disassembly of MPS is also one of the earliest signs of DAI.

## Discussion

In this study, with the assistance of CFD, we established an AoC system to study the mechanical stress-induced DAI as well as its underlying cellular cascades. Compatible with high-speed calcium imaging and super-resolution microscopy, this system enables the observation of instantaneous cellular responses triggered by mechanical stress. Using this system, we investigated the correlation between the intensity of mechanical stress and the extent of DAI, identified a delayed local axonal [Ca^2+^]i surge escorted rapid disruption of MPS during the early stage of DAI.

### Microfluidic models to investigate DAI

In the mechanically impacted brain, DAI triggers multiple secondary degenerations, such as Wallerian degeneration and excitotoxicity, which happen hours to days following the onset of the impact ^14,32^. When adapting *in vitro* models to investigate DAI, it is necessary to choose a suitable injury method pertinent to the damage process *in vivo*. Various microfluidic models have been developed to explore the degenerative cascades triggered by mechanical insults *in vitro*. In most of such models, axons are entirely disrupted (axotomy), including vacuum-assisted axotomy ^4,17^, laser-based axotomy ^38^ and chemical-based axonal injuries ^39^. However, these axotomy models, while being very useful to study the axonal regrowth following full axonal disruption, are not suitable for studying the mechanisms underlying DAI.

Moreover, several non-disruptive and stretch-based injury models were developed by exerting strain and shear force with sudden physical force or motion to cultured neurons ^13,21^. In these models, diffuse injuries were indeed induced by sudden stretching of axons on their axial direction without complete severing. However, as the whole neuronal cell body was also deformed by the stretch, damages were not restricted to the axon. As the somatodendritic region hosts the majority of organelles and machineries, which are required for both cellular metabolism and signaling, deformation of these core sections usually generate anterograde degeneration, thus initiating different cell death pathways from the retrograde degenerative signals underlying DAI ^40^.

Therefore, to more accurately mimick DAI *in vitro*, a new platform is required to restrict the non-disruptive injury exclusively to the axonal region. In this study, with the CFD-assisted design, we develop a new AoC injury device, which, by controlling the flow rate of injecting fluid, enables the accurate and exclusive fluid-induced DAI onto the axons growing in the 1 mm wide middle injury chamber. Using this device, we not only created the non-disruptive axonal injury signified by the formation of multiple focal axonal swellings (FAS), but also for the first time defined the positive correlation between the amplitude of external mechanical force exerted on axon surface and the extent of DAI. Moreover, we observed various subcellular injury responses in multiple axons simultaneously. Besides, with the current design of our AoC, it is also feasible to apply two different cellular treatments or seeding two different types of neurons in the two separated soma reservoirs, which doubles the experimental throughput, offering a versatile and simple platform for pharmacological screening.

### Primary cellular damages underlying the force-induced axonal injury

In early stages (0-120 min) following the mechanical stress, primary cellular damages such as DAI scattering in the white matters cannot be detected using non-invasive medical imaging methods ^1^. Nonetheless, during this narrow window, DAI triggers multiple secondary degenerative cascades, such as Wallerian degeneration and excitotoxicity ^14,32^. Secondary injuries are relatively easy to be detected. However, at the stage of secondary injury, synaptic connections or cell bodies of the neurons bearing DAI are irreversibly diminished, leading to unavoidable permanent functional loss ^33^. Therefore in order to prevent or mitigate further injuries of the secondary stage, it is critical to capture and target the earliest cellular changes underpinning the primary injury, which is marked by the formation of diffuse axonal varicosities or swellings caused by mechanical stress ^7^. Moreover, both our recent study and evidence from other labs have found that diffuse axonal swellings are not only dynamic but also, in some extent, reversible ^28,37^. These recent findings suggest that the primary stage of DAI might not be the inevitable dead end as previously expected. We therefore seek to use this *in vitro* AoC to study the cellular responses that happen instantaneously upon mechanical stress at various intensities, which are otherwise inaccessible using axial stretch or axotomy approaches.

As previously reported, as early as 15 min following the mechanical impacts, periodic blebs forming near the Ranvier nodes after DAI ^8^, the time window of 15 min is generally regarded as the time scale of primary injury. Then within 2 hours, the calcium influx will activate Ca^2+^-dependent enzymes, such as Caspases ^9^ and Calpains ^33^, marking the start of secondary injury. In less than 12 hours, these activated enzymes cleave multiple components of the cytoskeleton such as neurofilament ^10^, microtubule ^11^, F-actin ^41^ and spectrins ^9^, disrupting the axon structure and blocking the cargo trafficking^10^. These two phenomena signify the fixation of the secondary injury and would lead to degeneration of the injured axon. Using the AoC system, we revealed that a delayed [Ca^2+^]i increase correlates with the formation of focal swellings of axon (FAS) and both happen only seconds following the non-disruptive stress. Whereas a much higher [Ca^2+^]i surge was observed together with axotomy, if harsher mechanical impacts were applied.

To our surprise, using time-lapse SIM, we also found a much earlier disruption of the axonal cytoskeleton following DAI. Instead of hours following primary injury, MPS was abolished only several minutes following mechanical impacts. The loss of MPS is first observed in DAI, similar to the process of dynamic F-actin rings disassembly underlying the axon degeneration induced by trophic factor deprivation ^16,36^. These results suggest a more dynamic subcortical actomyosin network are involved in response to mechanical stress, indicating a potential role of subcortical actin rings in mediating primary axonal injury. Given the newly identified role of NM-II in both axon regeneration ^42^and structural stability ^37^, it might be a potential therapeutic target for DAI alleviation.

In summary, we developed an AoC platform to study cellular responses during the primary injury stage of DAI in cultured neurons. Using this system, we established the positive correlation between the strength of injected mechanical force and the extent of cellular injuries. The occurrence of delayed local axonal [Ca^2+^]i and focal axonal swellings was also discovered within the narrow time frame of primary injury. To our surprise, a much rapidly disassembled MPS underpinning DAI was also observed shortly following the mechanical injury.

## Material and Methods

### Microfluidic device fabrication

The multi-layer microfluidic device was fabricated using standard photolithography ^43^ and soft-lithography techniques ^44^ as previous described. Briefly, the microfluidic device was designed using AutoCAD software. The top and bottom pattern of the microfluidic device was printed on two Chrome Masks via a high-resolution direct laser writer (157 nm F2 laser exposure system). Then, a clean silicon wafer was spin-coated with SU-8 to a height of 5 μm, baked to enhance film adhesion on substrate, and exposed to UV light through a template mask patterned with the bottom layer of the microfluidic device. After a post-bake step, a second layer of SU-8 was spin-coated on the wafer to a height of 100 μm, and the process was repeated using a mask having the top layer of the device. The wafer was then heated to facilitate cross-linking of the UV-exposed resists, resolved with SU-8 developer, and cleaned before use. A soft lithography process was carried out to fabricate the polydimethylsiloxane (PDMS) microfluidic device. The PDMS mixture (10:1 base elastomer to curing agent) was poured onto the wafer and cured at 80 °C for an hour. The PDMS slab was removed from the wafer, trimmed to individual devices, and punched to create cell seeding ports. The PDMS slab was plasma treated, seeled against a glass-bottom dish, and immediately immersed in ethanol to enhance hydrophilicity of the PDMS surface.

### Neuronal culture and transfection

Under sterile conditions, glass-bottom dish attached with PDMS microfluidic device were washed with first 100% ethanol then Phosphate buffered saline (PBS) for 3 times before coated with 0.5 mg/mL Poly-L-Lysine in Borate buffer (1M, pH 8.5) in all chambers overnight. Hippocampal neurons were cultured from embryonic day 18 (E18) embryos of Sprague Dawley rats. All experiments were performed in accordance with relevant guidelines and regulations as approved by the University of Queensland Animal Ethics Committee (approval number: QBI/142/16/NHMRC/ARC). Hippocampal neurons were prepared as described previously ^45^ and were seeded into the soma chambers of microfluidic devices, briefly as follows: hippocampal neurons were dissociated and counted and a minimum of 5×106/mL concentrated cell suspensions were used for seeding purpose. Only soma chambers at both sides of the central injury channel were filled with above mentioned cell suspensions, with all spare suspensions removed from the reservoir. After the seeding step, chambers were placed in the CO2 incubator for 10-20 min to allow cell attachment, then culture medium (Neurobasal medium, 2% B27 and 1% L–GlutaMAX) was added to all reservoirs and axon chambers of the devices for neuronal growth. Then on 7 days in vitro (DIV7), hippocampal neurons in devices were transfected with indicated plasmids using Lipofectamine as previously described^45^. Microfluidic injury experiments were performed on DIV10.

### Confocal live-imaging of microfluidic-induced DAI

Live-imaging of microfluidic injury experiments were acquired with a Zeiss LSM710 inverted point-scanning laser confocal microscope, using 63 × 1.4 NA/190 μm WD/0.132 μm/pixel (1,024 × 1,024) objective in 37°C and 5% CO_2_. Imaging medium was Neurobasal minus phenol red (12348017; Thermo Fisher Scientific). Microfluidic injury was created by injecting the conditioned culture medium into the central injury channel by the Pump 11 Elite Programmable Syringe Pumps (Harvard Apparatus, #704505) at indicated flow rates for 3-5 min, while time-lapse images were acquired simultaneously. Diameter analysis of axons in time-lapse images were performed using ImageJ software (National Institutes of Health, v2.0.0/1.52p). Briefly, only the Lifact-GFP channel was selected for analysis, then images were set to 8-bit, background noise was filtered with 1 pixel Gaussian filter, converted to binary images by applying 6-255 threshold; Then separate region of interest (ROIs) were selected with 200×200 pixels size at the segments proximal to the channel exits. The minimum angle was measured by connecting the beginning and ending points of the axon segment using the straight-line tool. Five points with equal intervals along the axial length of axon segments were selected in each ROI. Diameter at these five points were measured using straight-line tool. Kymographs were generated using ImageJ software using the plugin Multi-Kymograph for ImageJ. All measurements were analysed using Graphic Prism, then all images and figures were compiled using Illustrator CS 23.1.1 (Adobe).

### Time-lapse SIM to detect MPS alternations underlying DAI

DIV10 rat hippocampal neurons cultured in microfluidic devices were subjected to live-labelling using SiR-actin, newly dissolved SiR-actin was added to culture medium at dilution rate of 1/10,000 followed by incubation for 1 h at 37°C CO_2_ incubator. These neurons in the microfluidic device were then washed once with warm phenol-red free Neurobasal medium before imaging at 30°C, using an Alpha Plan-Apochromat 100×/1.46 NA oil-immersion objective on a Zeiss ELYRA PS.1 SIM/PALM/STORM (SIM/photo-activated localization microscopy/stochastic optical reconstruction microscopy) super-resolution microscope (Carl Zeiss) built around an inverted Axio Observer.Z1 body and equipped with an sCMOS camera (PCO AG) for SIM acquisition and an iXon Ultra 897 electron multiplying charge-coupled device camera (Andor Technology) for PALM/STORM and controlled using ZEN 2.1 (Black) version 11.0. In the central injury chamber, the SiR-actin labelled neurons were imaged with the Fastframe mode (100-ms exposure time, a time series of 200 frames at 20-s intervals, a SIM grating size of 51 μm at 640-nm excitation and using five rotations). 2D structured illumination images were then reconstructed from the raw SIM data and channel alignment performed using Zen software. To ensure proper alignment of all channels, 4 channel SIM data were acquired and processed using 100-nm multi-spectral spheres mounted on a calibration slide (1783–455; Carl Zeiss) and channel alignment performed within Zen using the method Affine to provide a stretch and rotation dimension to the alignment, and the resulting data table was saved in a BIN file to be applied on multichannel specimen data.

### Numerical simulation of mechanical forces in microfluidic device

The FLUENT code in ANSYS software 18.0 was adapted to obtain the flow structure in this microfluidic device and force distribution on axon surfaces. A finite-volume method (FVM) was used to solve the steady three-dimensional Navier–Stokes (N–S) governing flow equations ^46,47^. The fluid was water with density of 998.2 kg/m^3^ and the viscosity of 0.001003 kg/m·s. The pressure field and the velocity field were coupled with the SIMPLE algorithm. The second order upwind was adapted to discretise the convection term, so does the diffusion term with the second-order central difference schemes. The inlet mass flow rate and the outlet pressure conditions are specified at the inlet and outlet. The non-slip and adiabatic wall boundary are imposed on the microchannel walls and axon surfaces. The double precision was empowered because of small dimension in microfluidic device. The simulation was performed on High-Performance Computer (HPC) with eight nodes. All the meshes are structural, and the mesh number with the reservoirs is about 16,000,000. It takes about three days to converge. The continuity equation, momentum equation, and energy equation are described as follows,

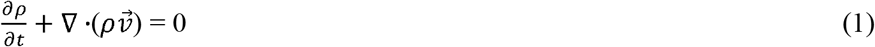

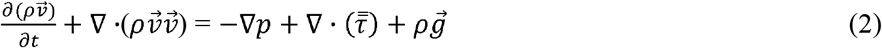

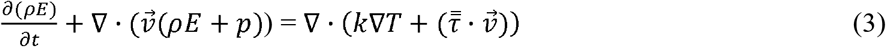

where 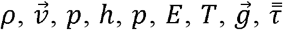 and λ are the density, velocity, static pressure, internal energy, static pressure, total energy, static temperature, gravitational acceleration, stress tensor and effective conductivity.

### Statistics

We used GraphPad Prism 7 (GraphPad Inc.) for statistical analyses. Results are reported as mean□±□s.e.m. For group comparisons, two-tailed nonparametric *t*-tests was executed. P values□<□0.05 indicated statistical significance. No statistical methods were used to predetermine sample sizes. Data distribution was assumed to be normal, but this was not formally tested. There was no formal randomization. Data collection and analysis were performed by different operators, who were blind to the conditions of the experiments.

## Supporting information

sMov. 1

sMov. 2

## Author Contribution

ZCX and WT designed the study. WL and WHF designed and made the microfluidic devices. WL, WHF and WT performed the live-imaging experiments. GD and DN performed the dissection and dissociation of rat neurons. PXR analysed the confocal data. WL and LZY analysed the numerical results. WL, WT, ZCX and WHF wrote the paper.

## Acknowledgement

We thank Frederic Meunier, Rumelor Amor, Tingting Tan and Jianfeng Wei and Rachel Gormal for their expert technical assistance. This work was supported by The Australian Research Council (grant DE170100546 to WT ant FT140100726 to CXZ), the National Natural Science Foundation of China (grant 31871036 to WT), Shanghaitech University start-up funds (to WT). The authors declare no competing financial interests. Author contributions: WT and CXZ designed the study, supervised the project, and wrote the manuscript. WT and WH performed live-imaging microscopy. LW and WH designed and fabricate the microfluidic devices. LW and LZ conducted the CFD simulation. PX analysed the live-imaging data. DN and GD performed the culture of primary rat hippocampal neurons. We acknowledge the supports from High Performance Computing of Queensland University of Technology. This work was performed in part at the Queensland node of the Australian National Fabrication Facility (ANFF-Q), a company established under the National Collaborative Research Infrastructure Strategy to provide nano- and micro-fabrication facilities for Australia’s researchers.

## Supplementary Material

**Supplementary Movie 1. Dynamic [Ca^2+^]i changes in axons subjected to axotomy induced by microfluidic injection.** Rat hippocampal neurons expressing GCaMP-6f were subjected to microfluidic-induced axotomy on DIV 10. Time-lapse images were acquired using open-field microscopy in the central injury chamber, at the injection flow rate of 1000 μL/min. Representative time-lapse image showing the disruptive process of transfected axons is shown, with [Ca^2+^]i level indicated by the fluorescent level of GCaMP-6f. Bar=5 μm.

**Supplementary Movie 2. Dynamic [Ca^2+^]i changes in axons subjected to non-disruptive diffuse injury induced by microfluidic injection.** Rat hippocampal neurons expressing GCaMP-6f were subjected to microfluidic-induced non-disruptive axon injury on DIV 10. Time-lapse images were acquired using open-field microscopy in the central injury chamber, the injection of 200 μL/min flow was indicated with yellow arrow. Representative time-lapse image showing the injury process of transfected axons is shown, with [Ca^2+^]i level indicated by the fluorescent level of GCaMP-6f. Bar=5 μm.

